# Embodied processing in whisker somatosensory cortex during exploratory behaviour in freely moving mice

**DOI:** 10.1101/2024.09.24.614719

**Authors:** Luka Gantar, Matthew A. Burgess, Neveen Mansour, Joaquín Rusco-Portabella, Alžbeta Námešná, David Gill, Isabella Harris, Patrycja Orlowska-Feuer, Aghileh S. Ebrahimi, Riccardo Storchi, Rasmus S. Petersen

## Abstract

Sensory systems have evolved to solve computational challenges that animals face during behaviour in their natural environments. To illuminate how sensory cortex operates under such conditions, we investigated the function of neurons in whisker-related Somatosensory Cortex (wS1) of freely moving mice, engaged in tactile exploratory behaviour. By recording neural activity from wS1 whilst tracking the mouse body in 3D, we found that wS1 neurons are substantially modulated by body state (configuration of individual body-parts and their derivatives), even in the absence of whisker afferent input. Most neurons were modulated by multiple dimensions of body state, with the most prominently encoded being the angle of the head to the body and locomotion speed. Overall, our data suggest that sensory cortex functions as an embodied representation, which integrates signals from its associated sense organ within a body schema.

## Introduction

Sensory function has been intensively studied using the classical paradigm where stimuli are delivered to the sense organ of an anaesthetised, immobilised animal and neuronal responses recorded ^1,2^. However, the brain’s sensory systems evolved to guide natural behaviour in freely moving animals ^3–5^ and a long-standing open question therefore concerns the extent to which sensory function in the classical paradigm generalises to that during natural free movement. To address this question requires investigation of sensory function in freely moving animals, able to express their natural repertoire of body states and movements^6–10^. Such experiments have only recently become possible due to advances in methods for the simultaneous cellular-resolution measurement of neural activity^11^ and precise tracking of posture in freely moving animals^12–19^. Our aim here was to investigate how whisker-related somatosensory cortex (wS1) operates during natural behaviour when animals are free to move.

There are indications that the operation of sensory systems in general, including wS1, during free movement might involve phenomena unanticipated from the classical anaesthetized paradigm. First, in naturally behaving, freely moving animals, the senses do not operate in isolation but rather as ‘perceptual systems’, the hallmark of which is that they operate in concert both with each other and with head/body movement ^20,21^. Humans coordinate eye, head and body movements during visually guided tasks^21^. Rats/mice coordinate head and whisker movements during tactile exploration^22–24^. This suggests the potential existence of circuits that integrate afferent sensory signals with information about body state. Second, the discovery, in behaving head-fixed animals, that locomotion speed modulates neuronal activity across the brain^25^, including the sensory cortices^26–28^ and that such modulation goes beyond arousal^29,30^, raises the question of whether more profound body-related modulation might occur when animals are able to express their full locomotor repertoire. Third, although little investigation of sensory cortex has been carried out on freely moving animals, there is evidence that body posture modulates neuronal activity both in sensory thalamus^31,32^ and in sensory cortical areas ^33,34^. However, how wS1 operates during natural behaviour is not known.

To address this issue, we set out to investigate what drives the activity of neurons in wS1 of freely moving mice, engaged in tactile exploratory behaviour. We find that wS1 neurons are sensitive not only to whisker-surface touch, but they are also substantially modulated by body state (configuration of head/body and its derivatives), even in the absence of afferent input from the whiskers. wS1 neurons are selective to, and collectively encode, multiple dimensions of body state. Overall, our data suggests that wS1 is an embodied representation, which integrates signals from its associated whisker sense organ within a body schema.

## Results

To investigate the function of wS1 whilst freely moving mice engage in tactile exploratory behaviour, we implanted Neuropixels probes into wS1 (n=6) (Supplementary Fig. 1) and introduced the mice to a dark, open-field arena (illuminated by IR light, 860 nm) containing an object (Fig. 1a). Mice are intrinsically motivated to explore novel objects^35,36^ and we encouraged exploration by replacing the object every 6 minutes. To measure body state (specifically, the configuration of the mouse’s head and spine), we imaged the mice with a system of four cameras (30 fps). We tracked 8 keypoints on the mouse’s body (snout, left/right ear, neck-base, tail-base, 3 along spine) and 3 keypoints on the housing of the microelectrode implant, in each of the four image planes (Fig. 1b and Supplementary Video S1). We then reconstructed the 3D coordinates of each keypoint using an algorithm that uses constraints from biomechanical relationships between body-parts to correct for occlusion errors (Methods). The outcome was a 24D (8 keypoints, 3 coordinates) description of mouse body configuration (‘pose’) in each video frame (Fig. 1c and Supplementary Video S2). To visualize the range of behaviours (meaning stereotypical poses or sub-second pose sequences) that the mice exhibit in the arena, we applied tSNE dimension reduction to 1 s pose sequences (Fig. 1d). Mice exhibited a broad range of natural behaviours - including forward locomotion, left/right turns, rearing, climbing, object investigation, immobility and grooming (Supplementary Video S3) - characteristic of rodent open field exploration^37^.

**Figure 1.**
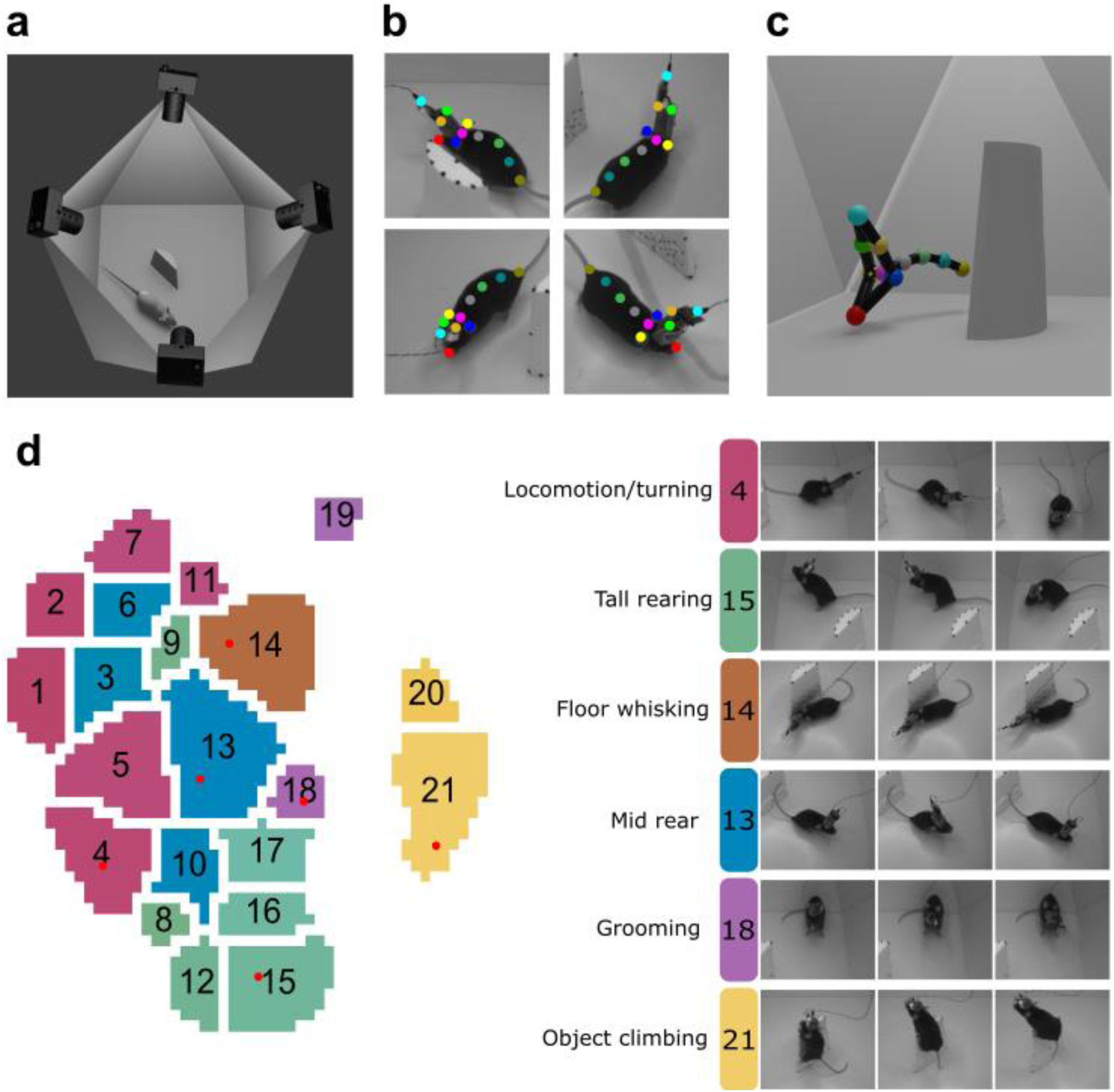
Tracking 3D body state of a freely moving mouse during open-field exploration. **a** Schematic of the behavioural arena, with camera locations indicated. An object was positioned in the middle of the arena. **b** Zoomed in view of a mouse from each camera, with keypoints tracked by DeepLabCut. **c** 3D reconstruction of the keypoints from the four 2D views in (**b**). **d** (Left) Behavioural clusters identified by t-SNE dimension reduction of 1 s pose sequences. Similar clusters were grouped, indicated by common colour. **d** (Right) Example clusters: each row shows the 1^st^, 10^th^ and 20^th^ frame of an example pose sequence. The location of each example in the tSNE plane is indicated by red dots in the left panel.

To investigate the relative role of tactile signals vs body state in driving the activity of neurons in wS1, we used a prediction paradigm^31,38–40^. We first aimed to estimate the extent to which wS1 firing rate can be predicted from whisker-surface touch. By tracking keypoints on the internal surfaces of the arena (object, walls and floor) and thereby reconstructing these surfaces in 3D, we calculated, in each frame, the distance from the mouse snout keypoint to the nearest point on the object, wall and floor (Fig. 2a): Snout Object Distance (SOD), Snout Wall Distance (SWD) and Snout Floor Distance (SFD) respectively. To construct a proxy for whisker-surface touch, we applied a compressive nonlinearity to the 3D vector [SOD, SWD, SFD] and refer to this as Snout Surface Distance (SSD) - (Fig. 2c and Methods). To estimate how well wS1 firing rate can be predicted from SSD, we trained a Supervised Learning algorithm (XGBoost) to predict the firing rate of each neuron (0.5 s bins) from SSD (N=436 neurons; Fig. 2b). Prediction accuracy was quantified by the fraction of firing rate variance explained (R^2^), which ranges from 0 to 1. Despite the complexity of the freely moving paradigm, we found that the firing rate of some neurons could be predicted with surprising accuracy (up to R^2^ = 0.64; Mean ± SD of R^2^ was 0.11 ± 0.11; Fig.2c-d) and that this was observed in every animal (range of mean R^2^ for individual mice 0.075-0.137; Fig. 2e). Predictions were statistically significant (R^2^ > mean + 3SD of shuffled distribution; Methods) for 92% of the neurons and this was again consistent across animals (individual mice 88%-97%; Fig. 2e). XGBoost predicted firing rate more accurately than a Generalised Linear Model (GLM) (R^2^ 0.11 ± 0.11 vs 0.06 ± 0.08), but the percentages of significant neurons were similar (Supplementary Fig. 2).

**Figure 2.**
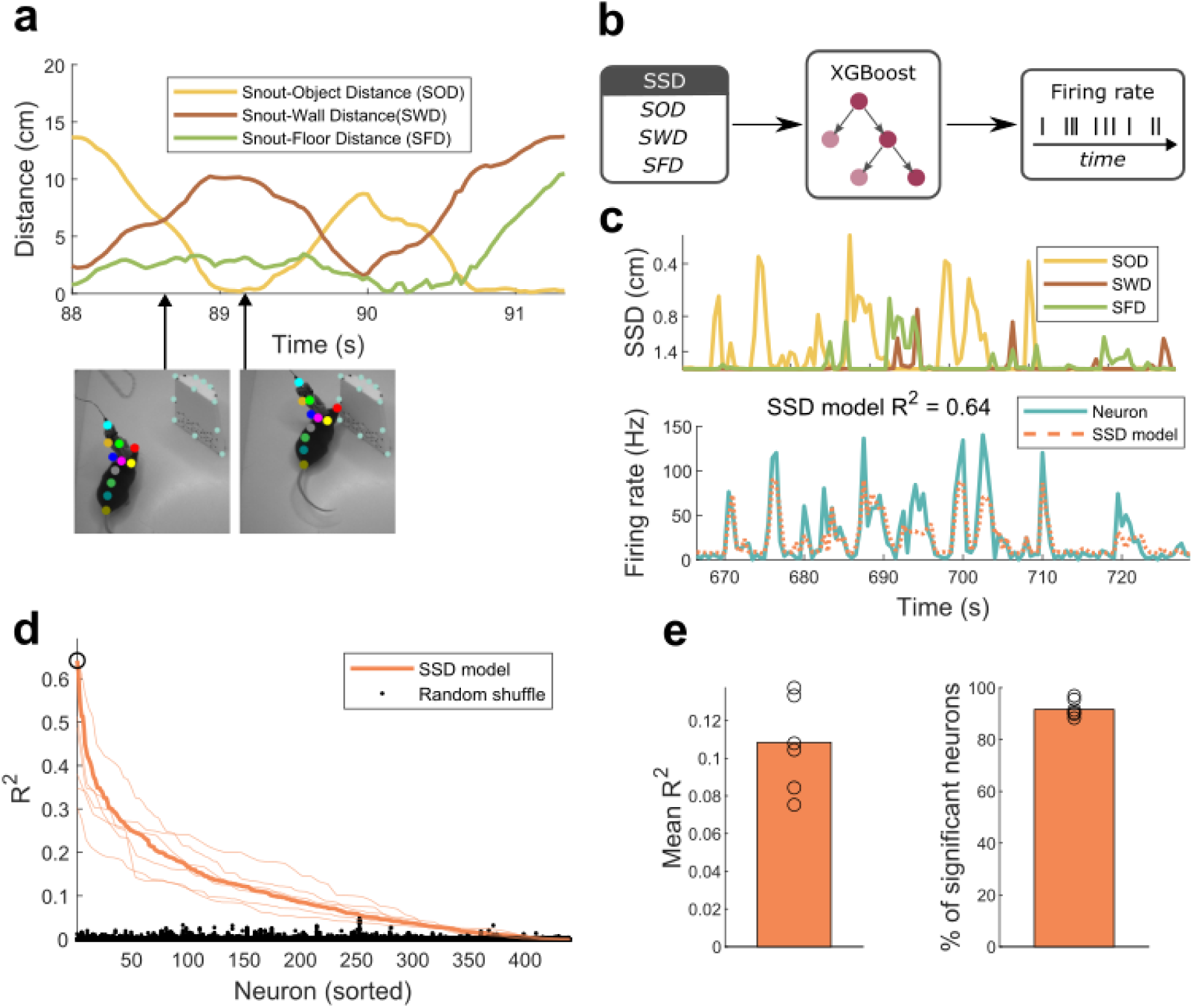
wS1 neurons encode whisker-surface touch. **a** Snout Surface Distance (SSD) during a 3 s episode in which a mouse explores an object. As the mouse approaches the object, it moves away from the wall; correspondingly, SOD decreases and SOW increases. Towards the end of the episode, the mouse rears against the object; here SOD remains low, but SFD increases. Camera images are shown for the two time points indicated by arrows, with tracked keypoints for both mouse and object superimposed. **b** Schematic of the XGBoost model used to predict neuronal firing rate from SSD. **c** Example of a wS1 neuron whose firing rate was well-predicted by SSD. Top: SSD is plotted on an inverse, nonlinear scale so that peaks correspond to touch events (low SSD) and SSDs beyond touch range saturate. Bottom: corresponding time series of the neuron’s firing rate along with that predicted by the SSD model. The firing rate peaks of the neuron correspond clearly to touch events (low SOD). **d** Variance explained by the SSD model (R^2^) for all recorded wS1 neurons (thick orange line) across 6 animals (thin orange lines); sorted by decreasing R^2^. Data for individual mice are rescaled to fit the x axis. R^2^ for the example neuron of panel **c** is denoted by the black circle. For each neuron, shuffled R^2^ values are also shown (black points, 20 shuffles per neuron). **e** (Left) Mean R^2^ across all neurons for the SSD model (orange bar) along with means for individual mice (black circles). (Right) Percentage of all neurons that significantly encode SSD (orange bar) along with values for individual mice (black circles). A neuron was deemed significant if its R^2^ exceeded mean + 3 SD of the shuffled distribution.

The next step was to investigate the extent to which wS1 neurons encode body state – that is, the spatial configuration of the 8 body keypoints (‘pose’) and changes in pose. 3D tracking provided a rich description of body state, capturing the 3D orientation of the animal (heading and rearing), its 3D location, 3D velocity, along with deformations in body shape (e.g., rotation of the head with respect to the body and extension-contraction of the body). We parameterized pose in terms of 3D head orientation (yaw, pitch, roll; Fig. 3a) and used a Statistical Shape Model^12,25–27,41^ to parameterize deformations of body shape in terms of their Principal Components (PCs). The first PC (‘PC1’) corresponded to changes in pitch angle of the head with respect to the body; PC2 to twist of the head with respect to the body and PC3 to extension/compression of the body (Supplementary Video S4). PCs 1-3 captured 85% of body shape variance. We parameterised pose change by the frame-to-frame differences in pose parameters (Δyaw, Δpitch, Δroll, ΔPC1, ΔPC2, ΔPC3) along with – to capture locomotion - frame-to-frame differences in mouse location (Δlocx, Δlocy, Δlocz) (Methods).

**Figure 3.**
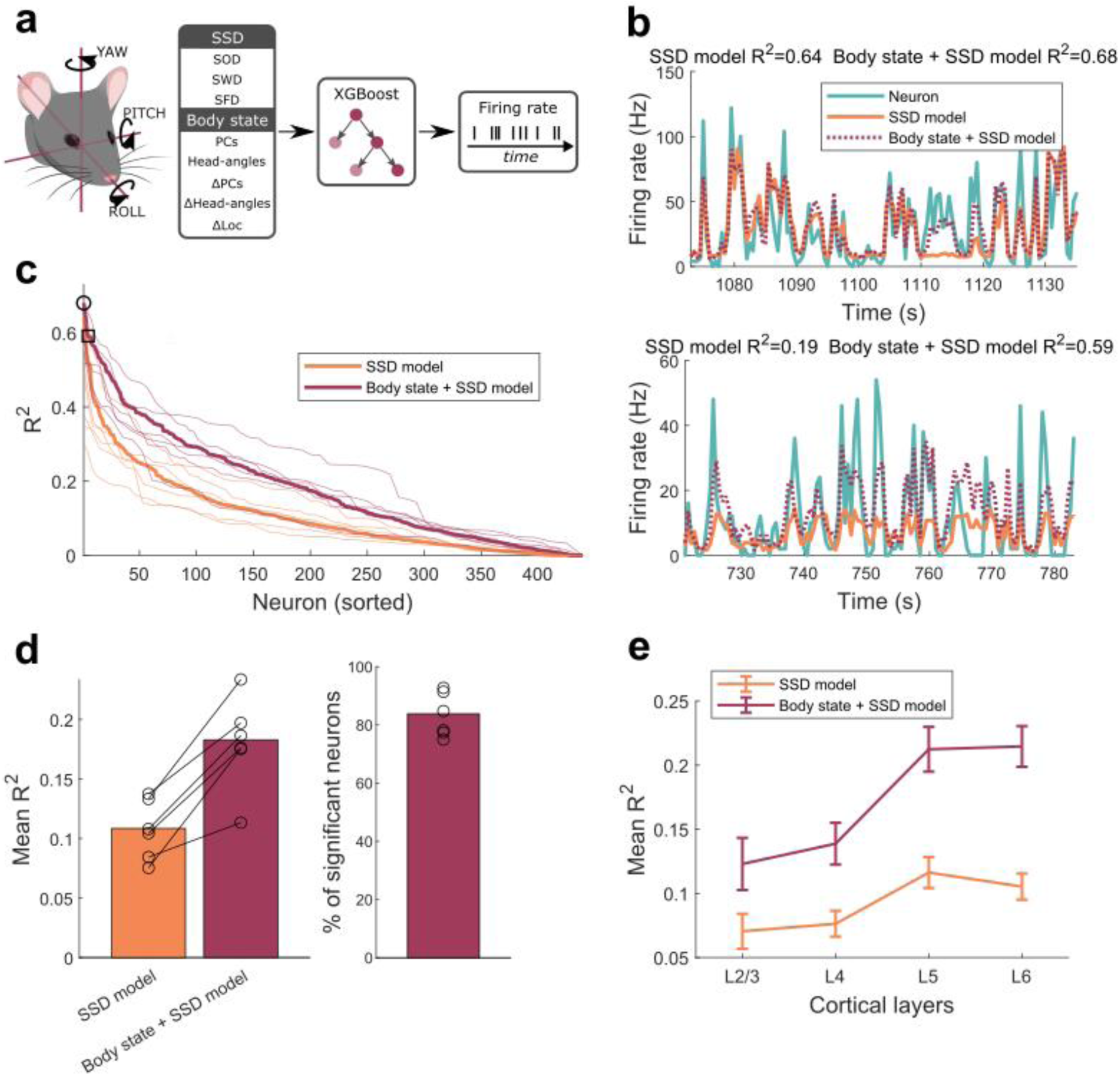
wS1 neurons encode body state. **a** (Left) 3D head angles. (Right) Schematic of the ‘body state + SSD’ Supervised Learning model used to predict neuronal activity. Parameters defined in main text. **b** Responses of two example neurons compared to those predicted by SSD model and body state + SSD model. **c** R^2^ for SSD model compared to that for body state + SSD model: data from all neurons (thick lines) and individual mice (thin lines). Neurons are sorted separately for each model. Neurons from (**b**) denoted by square and circle respectively. **d** (Left) Mean R^2^ for SSD model vs body state + SSD model for all data, with means for individual mice (black circles). (Right) Percentage of neurons encoding body state; defined as those where R^2^ for the body state + SSD model exceeds mean + 3 standard deviations of the R^2^ of a shuffled model trained on data where SSD is intact but body state is shuffled. **e** Mean R^2^ as a function of cortical layer for the two models (mean and SEM).

To determine the extent to which these body state parameters might be encoded by the activity of wS1 neurons, we trained XGBoost to predict firing rate from both touch (SSD) and body state parameters together (body state + SSD model) and compared it to the prediction performance based on SSD alone (SSD model). For some neurons, inclusion of body state parameters had minimal effect on prediction accuracy (Fig. 3b-upper). However, the more common finding was a substantial increase in prediction accuracy (Fig. 3b-lower; Fig. 3c). Overall, R^2^ for the neuronal population increased from 0.11 ± 0.11 to 0.18 ± 0.15 (Fig. 3d) - a median increase of 73%, and substantial increases were observed in all mice (individual mouse medians 37-138%; Fig. 3d). Overall, 84% of neurons showed a significant increase in prediction accuracy (individual mice 75-93%; Fig. 3d; Methods). Consistent results were obtained when using a GLM (Supplementary Fig. 3). There was a significant effect on prediction accuracy due to both model type (SSD, SSD+body state) and cortical layer (L2/3, L4, L5, L6) (2-way ANOVA; main effect of model type, p<0.001; main effect of layer, p<0.001). R^2^ for layers L5 and L6 were significantly higher than those for both L2/3 and L4 (p<0.01; Tukey post hoc test) but there was no significant difference either between L5 and L6 or between L2/3 and L4 (p>0.05).

It is possible that the previous analysis might over-estimate the role of body state on wS1 activity, if body state parameters were to correlate with aspects of touch not fully captured by SSD. Most obviously, self-generated touch occurs during grooming, and grooming is associated with a characteristic hunched posture. To eliminate this possibility, we removed whisker afferent input in a subset of mice (n=2) by bilateral section of the infraorbital nerve (ION). This procedure abolished the wS1 response to whisker stimulation (Fig. 4a) and we confirmed that this state was maintained for at least 8 days (Supplementary Fig. 4). If wS1 is driven by whisker touch alone, ION section should drastically reduce the ability to predict neural activity from body state. In contrast, if wS1 is driven by body state (in addition to touch), body state should maintain substantial predictive power. To test these predictions, we re-recorded wS1 activity 7 days after ION section as the mice explored the arena (N=123 neurons) and repeated the above analysis using a body state XGBoost model (Fig. 4b). We found that R^2^ for the body state model was high after ION section (0.14 ± 0.11), only slightly lower than R^2^ for the body state and SSD model prior to ION section (0.18 ± 0.13) (Fig. 4c-d). Again, a GLM produced consistent results (pre ION section R^2^ for body state + SSD model was 0.12 ± 0.10; post ION section R^2^ for body state model was 0.08 ± 0.08). Collectively, these data indicate that, during freely moving behaviour, wS1 neurons are substantially influenced by body state.

**Figure 4.**
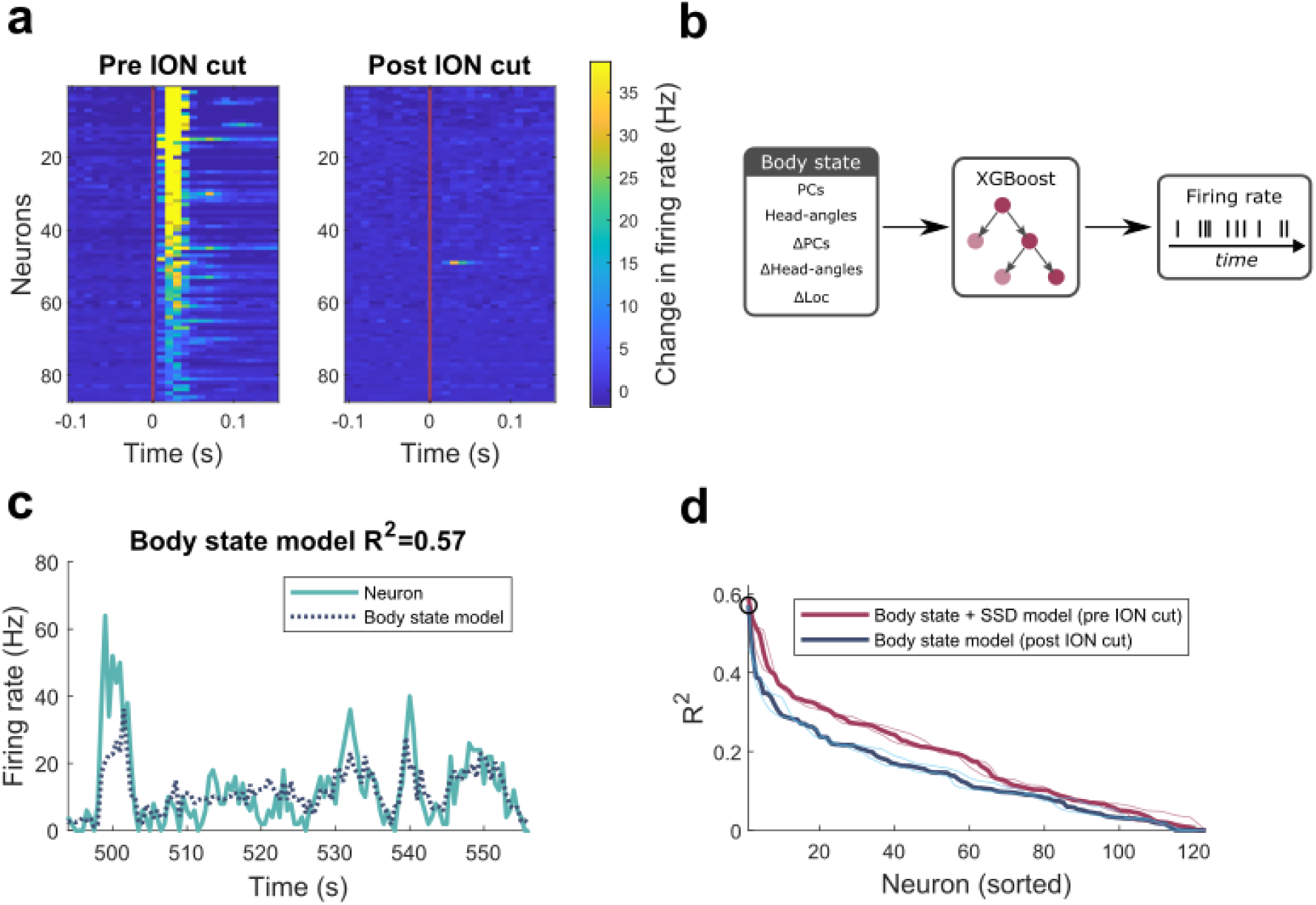
wS1 neurons continue to encode body state after removal of whisker tactile input. **a** Evoked response (change in firing rate compared to pre-stimulus firing rate) to whisker deflection delivered at time 0 (red vertical line), before and after ION section. Neurons are sorted by evoked response before ION cut and the same order is used in both panels. **b** Schematic of the body state model used to predict neuronal activity post ION cut. **c** Firing rate of an example neuron compared to that predicted from body state model. **d** R^2^ for body state model post ION cut in comparison to that of body state + SSD model pre ION cut: all data (thick lines); data from individual mice (thin lines). The neuron of panel (**c**) is denoted by black circle.

Which aspects of body state are encoded? To identify which out of the 15 body state parameters best predict firing rate, we trained a series of XGBoost models to predict firing rate from individual body state parameters in combination with SSD (N=234 neurons from 6 mice, where adding body state increased R^2^ by at least 0.05; Methods) and identified, for each neuron, its ‘best-predictor’ - the body state parameter that generated the highest R^2^. The neurons were diverse, but the two most common best-predictors were PC1 (pitch of head with respect to body; 37% of neurons) and Δlocx (forward locomotion velocity; 21%) (Fig. 5a). PC1 and Δlocx were also the most common parameters when this analysis was performed from data obtained after ION section (31% and 22% of neurons respectively; Supplementary Fig. 5a). Of the neurons whose activity was best-predicted by PC1, 23% preferred positive values of PC1 (‘head up neurons’), while 77% preferred negative values (‘head down neurons’) (Fig. 5b). Of the neurons best-predicted by forward locomotion, 75% responded to movement whilst 25% were suppressed (Fig. 5b).

**Figure 5.**
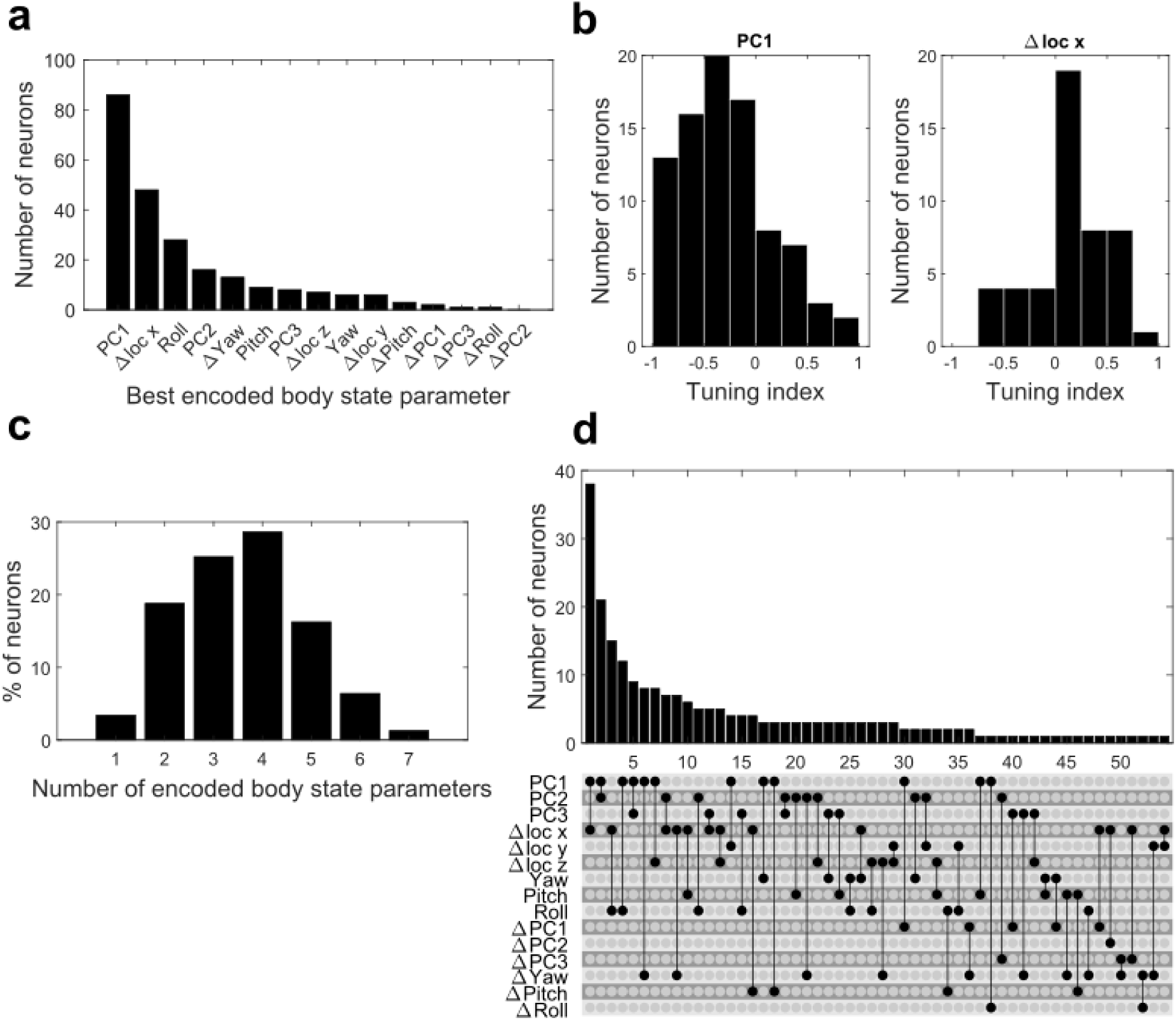
wS1 neurons encode multiple dimensions of body state. **a** Distribution of body state parameters that best-predicted neuronal activity (‘best-predictors’). **b** Histogram of tuning indices to PC1 and Δlocx respectively for neurons where these were the best-predictors. For each neuron, the tuning index for PC1 was the mean firing rate in frames where PC1 was high (>90^th^ percentile) minus the mean firing rate in frames where it was low (<10^th^ percentile), normalised by the sum of these two quantities. Thus, neurons that fires to high (positive) PC1 have positive tuning indices; those that fire to low (negative) PC1 have negative tuning indices. Tuning index for Δlocx defined analogously. **c** Distribution of how many body state parameters each neuron encodes (see main text for criterion). **d** Distribution of neuron types based on the pair of body state parameters that best predict their response. (Top) Frequency of observed neuron types. (Bottom) Combination of parameters that define a neuron type.

To determine whether neurons encoded single or multiple body state parameters, we used XGBoost to identify the body state parameter that added most predictive power to that already explained by the best-predictor and iterated this process until at least 90% of the variance explained by body state was accounted for (Methods). In this way, we found that 97% of neurons encoded at least 2 parameters (number of encoded parameters mean ± SD 3.6 ± 1.3; Fig. 5c) and this was also the case after ION section (mean ± SD 4.6 ± 1.5; Supplementary Fig 5c). Consistent with the previous analysis, the most common best-predicting pair was PC1 and Δlocx, but only 12% of neurons were of precisely this type (Fig. 5d): the neurons exhibited 54 unique combinations of encoded parameter pairs. Collectively, these data indicate a rich, multidimensional representation of body state in wS1.

## Discussion

The overall aim of this study was to investigate the function of sensory cortex during natural tactile exploratory behaviour when animals are free to move. Our main findings were that wS1 neurons are strongly modulated by body state, that this modulation is multidimensional and that it persists even in the complete absence of afferent whisker input.

Our findings reveal that sensory function during natural free movement has a pronounced integrative character, unanticipated from our knowledge of how wS1 operates in the classical experimental paradigm where animals are immobilised. As expected, wS1 neurons were sensitive to whisker touch but they were also strongly modulated by body state. One component of this modulation was forward locomotion: this is consistent with studies of wS1 in behaving head-fixed animals^25–28,42^, indicating that locomotion-modulation generalises to the freely moving state. However, our most striking finding was that modulation by body state is multidimensional and goes well beyond an effect just of forward locomotion. Indeed, the most prominent modulatory component was the pitch angle of the head to the body. The dimensionality of the body state modulation reported here should be taken as a lower bound: orofacial movement modulates dorsal cortex in head-fixed animals^29,30^ and tracking the body at higher resolution may reveal further dimensions of body-state modulation. Body state modulation was most pronounced in the infragranular layers. This may reflect the close anatomical association of the deep layers with the motor system^43–47^. Movement and planned movement modulate human S1^48,49^, indicating that body state modulation of S1 may be a general phenomenon.

Why might wS1 be modulated by body state? Active control of the sense organs is integral to perception^50–52^. When free to move their heads, rodents make coordinated movements of the whiskers and head^22–24^, and the activity of whisker low-threshold mechanoreceptors depends not only on the structure of the external surfaces being touched (exafference) but also on how the whiskers are moving^53^ and how the head /body is moving (reafference). Neural circuits that decode external structure from whisker afferent signals are thus likely to benefit from knowledge of head/body state. Our findings are consistent with the possibility that such integrative processing occurs already in wS1. A related perspective comes from normative modelling motivated by the concepts of Efficient/Predictive Coding^54–56^ which implies that neuronal receptive fields in, for example, V1, should depend on correlations across natural visual scenes^56–58^. In principle, the concept applies also across modalities. From the general standpoint of Efficient Coding, our basic finding of convergence of signals from whiskers and body state is expected from the behavioural phenomenon of whisker-body coordination.

What might be the nature and origin of the wS1 body state signal? Body state information may, in principle, reflect ascending drive from the proprioceptive/vestibular systems or efference copy from motor cortex. Considering first proprioceptive vs vestibular mechanisms, we found the body state modulation effect to be best-predicted in some neurons by allocentric head angles (roll, pitch, yaw and their derivatives) but more commonly by egocentric head angles (PC1 and PC2); in particular, PC1 (pitch angle of the head with respect to the body axis). Although caution is necessary in interpreting these data due to the various body state parameters being correlated, coding of allocentric parameters is most simply explained by a vestibular mechanism and that of egocentric parameters by a proprioceptive one. The preponderance of egocentric coding in our data thus tentatively points to a proprioceptive mechanism. Consistent with this interpretation, in our experiment, all head movements were self-generated and vestibular-sensitive neurons in VP thalamus are known to respond less to these movements than to kinematically-matched external ones ^59–61^). Conversely, during self-generated movements, proprioceptive input at spinal level is facilitated^62^.The proprioceptive ascending pathway projects via the ventromedial shell of VPL/VPM thalamus to S1^63,64^. Long-range corticocortical connections^65,66^ offer a potential substrate for proprioceptive input to the S1 dysgranular zone^67,68^ to reach wS1. Another possible explanation for our findings is that efference copy of movement commands in M1 is transmitted to wS1. Consistent with this possibility is the major projection from M1 to wS1 layer 5^46^ and our finding that body state modulation is strongest in the infragranular layers. Overall, proprioceptive and corollary discharge appear to be the most likely mechanisms, but further work is needed to resolve this.

In conclusion, this study provides a new appreciation of how the brain’s tactile system operates during natural exploratory behaviour. wS1 is an embodied representation, which integrates signals from its associated whisker sense organ with a rich representation of body state.

## Methods

### Animals

In this study, we used 6 female C57BL/6 mice obtained from Charles River or Envigo. All mice were initially stored in cages of four and housed individually after surgical implantation of the chronic electrode. Animals were provided with food and water *ad libitum* and kept on a 12:12 light-dark cycle. Mice were 10 - 13 weeks old, and weighed 18-21 g, on the day of the chronic implant surgery.

### Ethical statement

This study was carried out in accordance with the UK Animals Scientific Procedures Act 1986 and were approved by the University of Manchester’s Animal Welfare and Ethical Review Body (AWERB).

### Chronic implant surgery

Mice were anaesthetised with isoflurane (2% at 2 L/min O_2_) and head-fixed in a stereotaxic frame (Narishige SR-5N). Anaesthesia and a body temperature of 37°C were maintained for the whole duration of the surgery. Local anaesthesic (1% EMLA cream) was topically applied to the area surrounding the incision and analgesic administered (Buprecare, 0.05 mg/kg s.c.)). Drying of the eyes was prevented by regular application of eye gel (Lubrithal). Two screws (2.5 mm AP, 1.5 mm ML and -2.0 mm AP, -2.0 mm ML) and a grounding pin (WPI 5483; 2.5 mm AP, -1.0 mm ML) were implanted following craniotomies (1-2 mm) centred at their respective coordinates. The grounding pin and two skull screws were cemented in place using dental cement (RelyX™ Unicem 2 Automix) and cured with blue light (Starlight Pro; mectron). An additional craniotomy (1-2 mm) centred on probe implant coordinates: 1.5 mm AP, 3.1 mm ML was performed. DiI (Abcam) was carefully applied to the Neuropixels 1.0 probe (imec) for post-mortem histology and probe alignment, after which the probe was lowered into the brain at a speed of ∼10 μm/s (tip depth 3.76 mm beneath cortical surface). A ground wire was attached to the grounding pin and the probe tip was set as the reference site. The craniotomy was covered with Kwik-Cast (WPI) and the probe cemented both to the skull (RelyX™ Unicem 2 Automix), and to the plastic protective enclosure positioned around the probe^69^.

After the surgery, mice were individually housed and allowed to recover for 5 days. During this time, mouse behaviour and weight were monitored. Thereafter, a headstage (imec) was connected to the probe’s flex cable under brief anaesthesia (isoflurane, 2% at 2 L/min O_2_). Electrophysiological recordings began the following day.

### Electrophysiological recordings

The experimental setup was adapted from a Storchi *et al*.^70^. Briefly, the mouse was placed inside an open-field arena (width = 28 cm, length = 28 cm and height = 30 cm) made from non-reflective white Perspex and located inside a Faraday cage. Four cameras (Chameleon3 monochrome; FLIR) were mounted at each of the four corners of the arena. Video data (30 fps) was recorded (FlyCapture Software Development KIT; FLIR), and the cameras synchronised via an Arduino Uno Rev3 running a custom Python script. Recordings were performed in the dark (infrared illumination, ring light 860 nm, Stemmer Imaging) with cameras mounted with infrared low pass filter (720 nm, Edmund Optics, #65-796).

A (daily) recording session consisted of seven trials. Each trial lasted 6 minutes with a 3-minute inter-trial interval. The first trial was to habituate the mouse to the empty arena: no object was present. Prior to the start of the following six trials, one of a set of six objects was placed in the centre of the arena. The objects had the same half-truncated cone shape but differed in the relief on the front side. At the end of each trial, the object was removed and cleaned (70 % ethanol) and replaced. Object order was randomised on each recording day. At the end of each recording session, the animal was returned to its home cage, and the arena cleaned (70 % ethanol).

To verify probe location, animals were anaesthetised (sleep-mix) and responses to piezoelectric whisker stimulation recorded as above.

Extracellular potentials were recorded with Open Ephys^71^ (version 0.5.3 or 0.6.6). Both the camera triggers and electrophysiological data were acquired (PXIe Acquisition Module; imec)) at 30 kHz. Off- line, data from all 7 trials on a given day were first concatenated and then spike sorted with Kilosort4^72^. Clusters were accepted as well-isolated units if they met standard curation criteria based on refractory period violation rate of <0.1, waveform, firing rate and amplitude stability over time (Phy2).

### Infraorbital nerve cut surgery

Mice were anaesthetised with isoflurane (2% at 2 L/min O_2_) or with “sleep-mix”^73^ (fentanyl (0.05 mg/kg), midazolam (5.0 mg/kg), medetomidine (0.5 mg/kg) in saline solution (0.9%, i.p.)). The mouse was head-fixed in a stereotaxic frame as above and local anaesthesia, analgesic and eye gel were administered as above. An incision was made on the snout from the midpoint between the eyes to the nose tip. The skin was retracted and connective tissue separated by blunt dissection to expose the infraorbital nerve (ION). The nerve was then cut at the infraorbital foramen. The incision was sealed using tissue adhesive (3M 1469 Vetbond) and sutures (6/0 11mm Ethicon Mersilk). At the end of the procedure, the “sleep-mix” anaesthesia was reversed by administration of “wake-mix”^73^ (naloxone (1.2 mg/kg), flumazenil (0.5 mg/kg), atipamezole (2.5 mg/kg) in saline solution (0.9%; i.p.)). Whiskers were stimulated before and after the ION cut to confirm that the sectioning had removed afferent whisker input to the brain. Whiskers were inserted into a mesh and stimulated via a piezoelectric actuator (P/N PL127.10; Physik Instrumente). Using a function generator (RS PRO AFG21005), repeated square wave pulses (1Hz frequency with 50% duty cycle) were generated, amplified (E-650.00 LVPZT-AMPLIFIER, Physik Instrumente) and applied to the actuator and used to deflect the whiskers in the rostro-caudal direction. Extracellular potentials were recorded from the chronic implant as detailed above. After surgery, ION sectioned mice were individually housed and allowed to recover for 7 days, during which behaviour and weight were carefully monitored.

### Histology

Once all electrophysiological recordings were complete, mice were terminally anaesthetised with isoflurane (2% at 2 L/min O_2_) or urethane (1.6 g/kg, IP injection) and transcardially perfused with 20 ml of phosphate buffered saline (PBS, Sigma-Aldrich) followed by 20 ml of 4 % paraformaldehyde (PFA, Sigma-Aldrich) in 0.1 M PBS (pH = 7.4); both solutions kept at 4°C. The mouse brain was extracted and the implant carefully removed. Extracted brains were post-fixed for ∼6 h at 4°C and transferred to 30 % sucrose in 0.1 M PBS solution for ∼48 h until they sank. Brains were cut into 100 µm coronal sections with a freezing microtome (Leitz sledge microtome 1400 with Bright solid-state freezer) and stained for Cytochrome Oxidase (Wong-Riley, 1979).

Brain sections were imaged with a fluorescence microscope (Olympus BX63 single slide scanner) under brightfield and fluorescence with Texas-Red filters to visualise the section structure and probe track respectively.

To validate that the probe was implanted in wS1, sections containing the probe track were aligned to the Allen Brain Atlas (MATLAB, https://github.com/petersaj/AP_histology). Localisation of the probe in wS1 was also verified by checking for the presence of whisker-evoked responses in the recordings carried out under anaesthesia (see ‘Electrophysiological recordings’). To estimate the sub-pial depth of the spike-sorted units, we first identified the depth of each unit in probe coordinates (relative to the probe tip). Next, we subtracted the probe depth corresponding to the cortical surface (identified from recorded signals) to yield sub-surface depth. The laminar depth was assigned using laminar boundaries from the aligned atlas section. As a consistency check, we verified that the sub-surface unit depths corresponded to the region between the cortical surface and the border of the subcortical white matter.

### 3D mouse tracking

DeepLabCut^13^ (version 2.2.2) was used to track 11 keypoints on the mouse body (snout, left ear, right ear, three points on the implant, neck base, body midpoint, tail base, neck-body midpoint and body-tail midpoint) in the video from each of the four cameras. 2158 frames were extracted (k-means clustering in DeepLabCut) across all cameras and mice. Keypoints on these frames were manually labelled and used to train a Deep Neural Network (ResNet-101). To enable 3D reconstruction of the keypoints, the cameras were calibrated^70^, defining an arena coordinate system (x and y axes defining the horizontal plane, z the vertical).

The next step was to reconstruct (‘triangulate’) 3D keypoint coordinates from the 2D keypoint estimates in the four cameras. The standard Direct Linear Transform^74^ typically worked well in frames where a given keypoint was accurately tracked in each of the four views. However, where the body part corresponding to a given keypoint was occluded in one or more views, this approach generated errors. To address this, we used a robust Maximum A Posteriori (MAP) approach. Due to biomechanical constraints, there are strong statistical relationships between keypoints on the head and body and we used this as prior knowledge to constrain the reconstruction. Both likelihood (*P_L_*) and prior (*P_P_*) were approximated as multivariate Gaussians. The likelihood was a product over camera-specific likelihoods *P_L_(data_i_|x)*, where *i* = 1…4 indexes the cameras, *data_i_* is the video data for a given frame in camera *i* and *x = [x_1_,x_2_,…]* is the concatenated vector of 3D coordinates for keypoints *x_j_* in that frame. The mean of the *P_L_(data_i_|x)* was estimated by applying the pseudoinverse of the *i*th camera calibration matrix to each 2D keypoint estimate from the *i*th camera. The likelihoods for the keypoints were approximated as uncorrelated. The reliability (inverse covariance) of *P_L_(data_i_|x)* for each keypoint in the image plane of camera *i* was taken as inversely proportional to the DeepLabCut confidence value for that keypoint. Reliability orthogonal to the image plane was set to zero. Maximum likelihood estimates for *x* were calculated using the analytic solution for a Gaussian ^75^ and the set of these estimates used to estimate the mean and covariance of the prior *P_P_*. Finally, MAP solutions for *x* were calculated using the analytic solution (Gaussian case) for the maximum of the Posterior *P_L_P_P_*.

A post-processing step was applied to frames where the snout-ear distance were outliers (outside the range of 2^nd^ – 98^th^ percentiles). To obtain a refined snout keypoint estimate for such an outlier frame, we first calculated the median distance from the snout to four other keypoints (left ear, right ear, neck base, top of implant housing) across the dataset. By numerical optimisation, we then found the coordinate whose distance to each of these four keypoints in the outlier frame best matched the median distances.

### t-SNE analysis

To visualise the range of behaviours (by which we mean mouse pose and pose trajectory over a short time period) that the mice exhibited during the above chronic recordings, we parsed the 3D keypoint data from each mouse into 1 s (30 frame) chunks. To remove variance due to the mouse’s starting location and starting orientation, we first normalised the chunks so that the mouse’s location and orientation in the first frame of each chunk was the same. Specifically, for each chunk, we used the Procrustes algorithm (Matlab implementation) to identify the Euclidean transformation that aligned the pose in the first frame to a common reference pose and then applied this transformation to each frame in the chunk. These chunks were 720D (8 body keypoints, 3 coordinates, 30 frames) but 99% of the variance was explained by the first 20 Principal Components (PCs). To obtain an ethogram, t-SNE was applied to the first 20 PCs, followed by smoothing with kernel density estimation (bandwidth [3, 3]) and watershed clustering^76^.

### Extraction of Snout Surface Distance and body state parameters

We constructed a proxy for whisker-surface touch based on the distance of mouse’s snout to the nearest touchable surface (object, walls and floor). To obtain Snout Object Distance (SOD) and Snout Wall Distance (SWD), we manually labelled keypoints on the object, floor and walls in images from each of the four cameras, and constructed 3D points as detailed above. Using these points, we calculated a dense grid of points over the walls, floor and exterior object surface. Then, in each frame, we measured Snout Object Distance (SOD) and Snout Wall Distance (SWD) as the minimum Euclidean distance from the snout keypoint to the object and wall points respectively. To obtain the Snout Floor Distance (SFD), the vertical distance coordinate of the snout keypoint from the floor was used. Since whiskers can only touch surfaces close to the snout, the final step was to pass these distance measures through a compressive nonlinear function 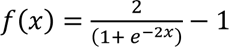 that saturates at *x* ∼ 2cm. We refer to this quantity [f(SOD,f(SWD),f(SFD)] as Snout Surface Distance (SSD).

To parameterise the mouse’s body pose in a given frame, we first aligned the mouse’s 3D pose (the set of 8 3D keypoint coordinates on the body) to a reference pose, by a Euclidean transformation in the horizontal plane (Procrustes algorithm). Residual variance in the aligned poses largely reflected the effects of non-rigid changes in pose (e.g., head turns, rearing). To capture these dimensions of body shape, we applied PCA to the aligned poses (using data from all mice). As reported in results, the first three PCs explained 85% of the total aligned-pose variance and were thus used for subsequent analysis. Allocentric head angles (Yaw, Pitch and Roll) were obtained by identifying an orthonormal basis describing the orientation of the mouse’s head (plane defined by the snout and ear keypoints, and the axis orthogonal to this plane) and calculating the 3D rotation needed to align the basis to a fixed reference basis (Procrustes algorithm). To extract whole body locomotor movements, we calculated the change in centre of mass of a mouse in a mouse-centred coordinate system where *x* is the direction from tail-base to snout (Δlocx); *y* the direction from ear to ear (Δlocy) and *z* is orthogonal to the plane defined by *x* and *y* directions (Δlocz). Specifically, the pose at each frame *n* was first aligned to the reference pose by a Euclidean transform (Procrustes algorithm). This same transformation was then applied to the pose of frame *n*+1 and the centres of mass of the two (transformed) poses calculated. In this coordinate system, change in the *x*, *y* and *z* coordinates correspond to forward location, turning and climbing/rearing respectively.

All the obtained motor parameters were filtered with a Savitzky-Golay filter (order 3 and frame length of 15). The derivatives of PCs and head angles were then calculated to create the final six motor parameters used.

### Firing rate prediction

We trained Supervised Learning models to predict neuronal firing rate from one or more inputs (bin size 0.5 s). For the ‘SSD’ model, the inputs were SOD, SWD and SFD. For the ‘body state’ model, the inputs were the 15 parameters of body state defined above. We used XGBoost and Poisson Generalized Linear Models (GLM) using xgboost 1.7.3 and pyglmnet 1.1 Python libraries, respectively, in Python 3.10. Data was randomly split into two equally sized sets. For each neuron, models were then trained on each set, and tested using the held-out set. Performance of a given model was quantified as the fraction of a neuron’s firing rate variance explained by the model (R^2^). In rare cases where the model’s predictions were less accurate than the mean of the firing rate, R^2^ was set to 0.

To determine if a neuron was significantly modulated by SSD, we compared the model’s R^2^ to that expected by chance (R^2^_null_). R^2^_null_ was calculated for each neuron by shifting SSD values randomly in time (18-180 seconds) with respect to the neuron’s spike train and training the model on this shifted data. This process was repeated 20 times for each neuron, and the resulting R^2^ values used to estimate the mean and variance of that neuron’s R^2^_null_ distribution. A neuron was deemed significantly modulated by SSD if its R^2^ exceeded the mean + 3 SD of its R^2^_null_ distribution. Similarly, to test whether one or more parameters (e.g., SSD) modulated neuronal activity to an extent greater than could be explained by another set of parameters (e.g., body state), a null distribution was obtained by selectively shifting the body-state inputs and then proceeding as above.

To evaluate the role of individual body state parameters, we trained a set of 15 XGBoost models to predict firing rate from the SSD parameters in combination with one of the body state parameters. The body state parameter associated with the largest significant R^2^ (the ‘best predictor’) was thence identified. The model consisting of the SSD parameters and the best predicting body state parameter was then used as a baseline and process repeated to identify the next-best predictor (that is, the body-state parameter that increases R^2^ most in comparison to the baseline model), and so on. Only neurons whose ‘body state + SSD’ R^2^ exceeded that of the SSD by 0.05 were included in the analysis of Fig. 5. The number of encoded body state parameters for a neuron was defined as the number of parameters necessary to account for 90% of the R^2^ explained by that neuron’s full model (SSD and all body state parameters).

To determine the tuning index of PC1 and Δlocx, we first determined the top and bottom deciles of the specific body state parameter for each mouse. We then calculated the average firing rate of each neuron during periods when the body state parameter was in the top and bottom deciles. The tuning index was then defined as the difference between these firing rates normalised by their sum.

## Acknowledgements

We thank Dario Campagner and Miguel Maravall for discussion; Ben Efron and Inbar Saraf-Sinik for assisting with training and protocol development; the University of Manchester Biological Services Facility and Bioimaging Facility for expert technical support. This study was funded by the Biotechnology and Biological Sciences Research Council (BB/V009680/1) and a Weizmann UK Making Connections Grant. The funders had no role in study design, data collection and analysis, decision to publish, or preparation of the manuscript.

## Contributions

LG, MAB, and NM contributed equally to this work. Conceived the project: RSP, RS, LG, MAB; funding acquisition: RSP, RS; methodology: RS, MAB, LG, PO-F, NM, AN, JR-P, ASE; data collection: MAB, NM, LG, IH; data analysis: LG, MAB, DG; software: LG, RS, RSP, ASE; training/supervision: MAB, RS, RSP, PO-F; writing (original draft): RSP, LG, NM; writing (editing): RSP, RS, LG, MAB.

