## Supplementary figures for "Embodied processing in whisker somatosensory cortex during exploratory behaviour in freely moving mice"

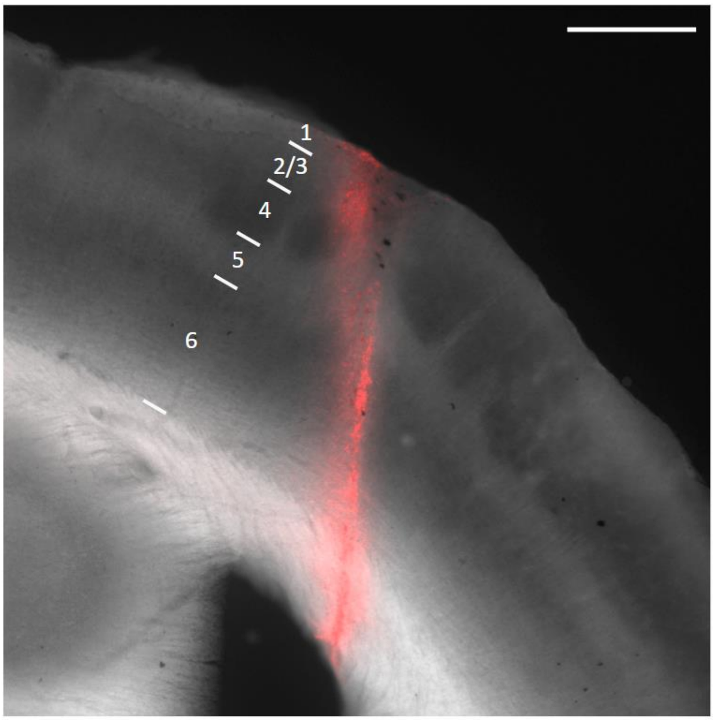


**Supplementary Figure 1. Microelectrode implant in wS1**. Prior to implant, the probe was coated with DiI (Methods). Cytochrome Oxidase stained section with Texas red channel (red) to show the DiI. Layer boundaries indicated (Methods). Scale bar is 500 µm.

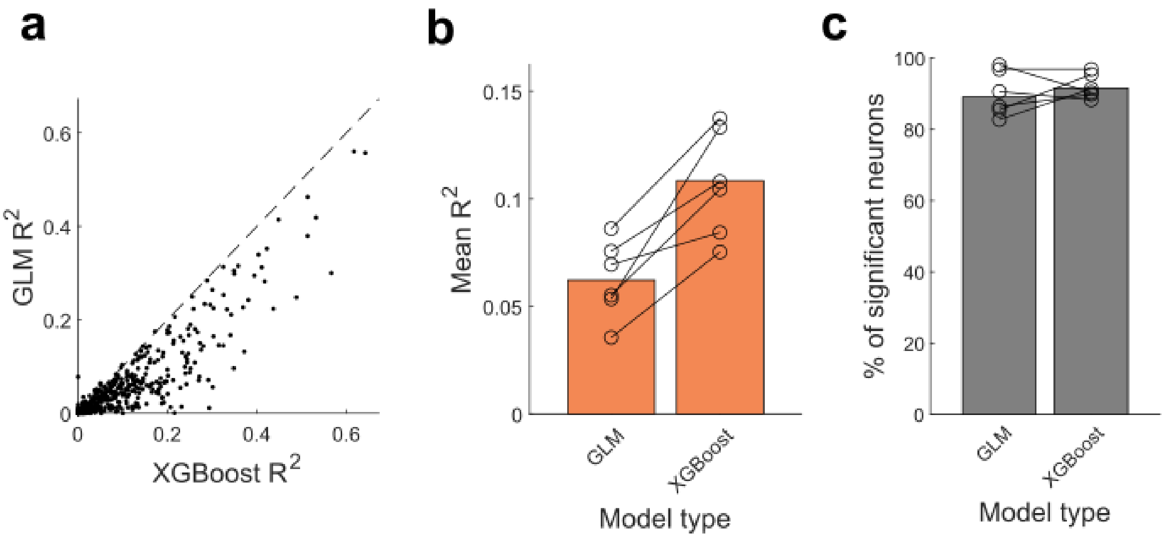


**Supplementary Figure 2. XGBoost predicts wS1 activity from Snout Surface Distance (SSD) more accurately than a Poisson GLM. a** Firing rate variance explained ($R^2$) by XGBoost compared to a Poisson GLM $R^2$ when the models were trained to predict firing rate from SSD: each dot one neuron. Same data set as Fig. 2. **b** Mean $R^2$ for all recorded neurons (orange bars) across 6 animals (black circles) **c** Percentage of neurons significantly encoding SSD with individual mice values as black circles (evaluated as in Fig. 2e).

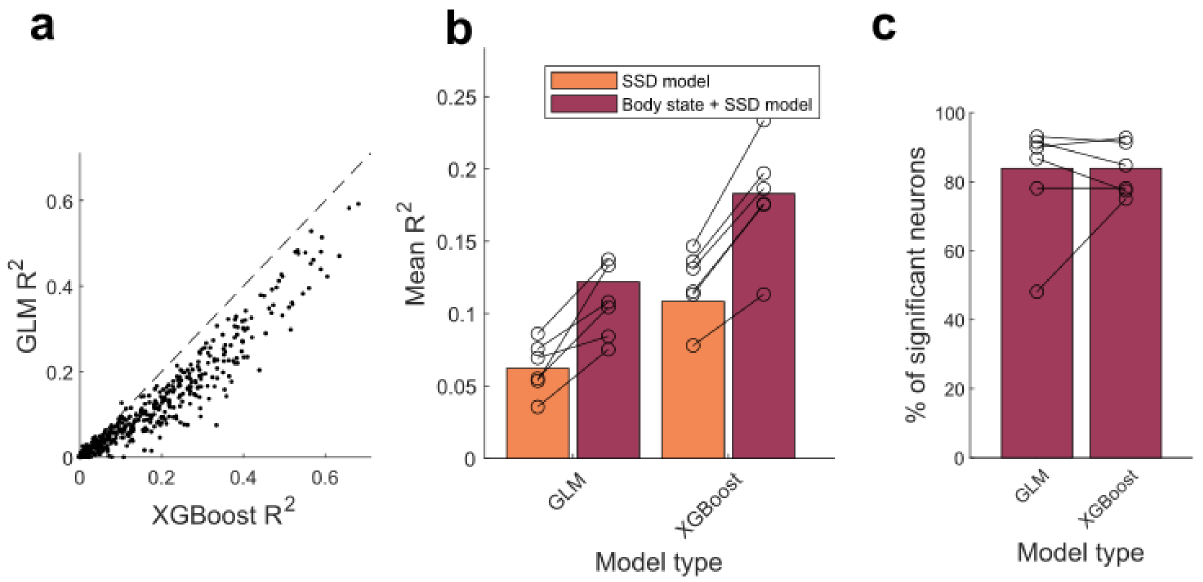


**Supplementary Figure 3. wS1 activity is predicted more accurately when body state descriptors are included, both with XGBoost and Poisson GLM models. a** Firing rate variance explained ($R^2$) by XGBoost compared to a Poisson GLM $R^2$ when the models were trained to predict firing rate from body state and SSD: each dot one neuron; same data set as Fig. 3. **b** Mean $R^2$ for all recorded neurons (orange/burgundy bars) across 6 animals (black circles). **c** Percentage of neurons significantly encoding body state with individual mice values as black circles (evaluated as in Fig. 3d).

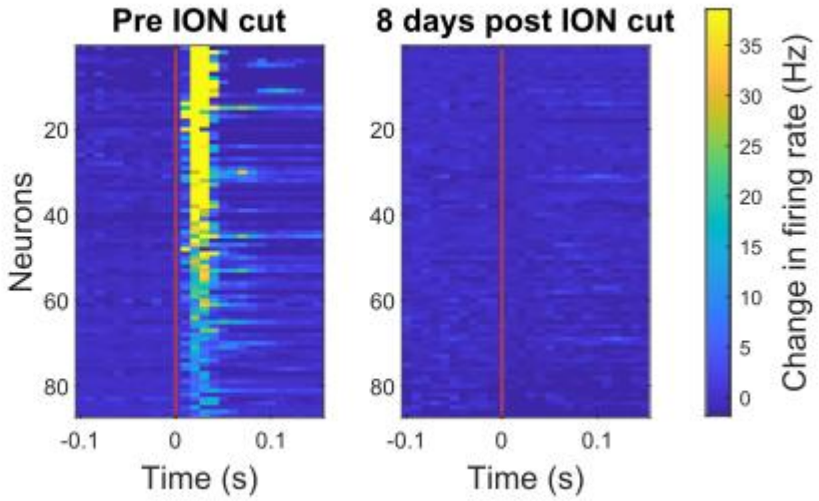


**Supplementary Figure 4. Whisker deafferentation is maintained at least 8 days after ION section.** Evoked response (change in firing rate compared to pre-stimulus firing rate) to whisker deflection delivered at time 0 (red vertical line), immediately before, and 8 days after ION section. The vertical red line indicates the time of the whisker deflection. In left panel (repeated from Fig. 4 for convenience), neurons are sorted by evoked response. It is unlikely that the same neurons were recorded after 8 days: the rows in left and right panels should not be assumed to correspond to the same neuron.

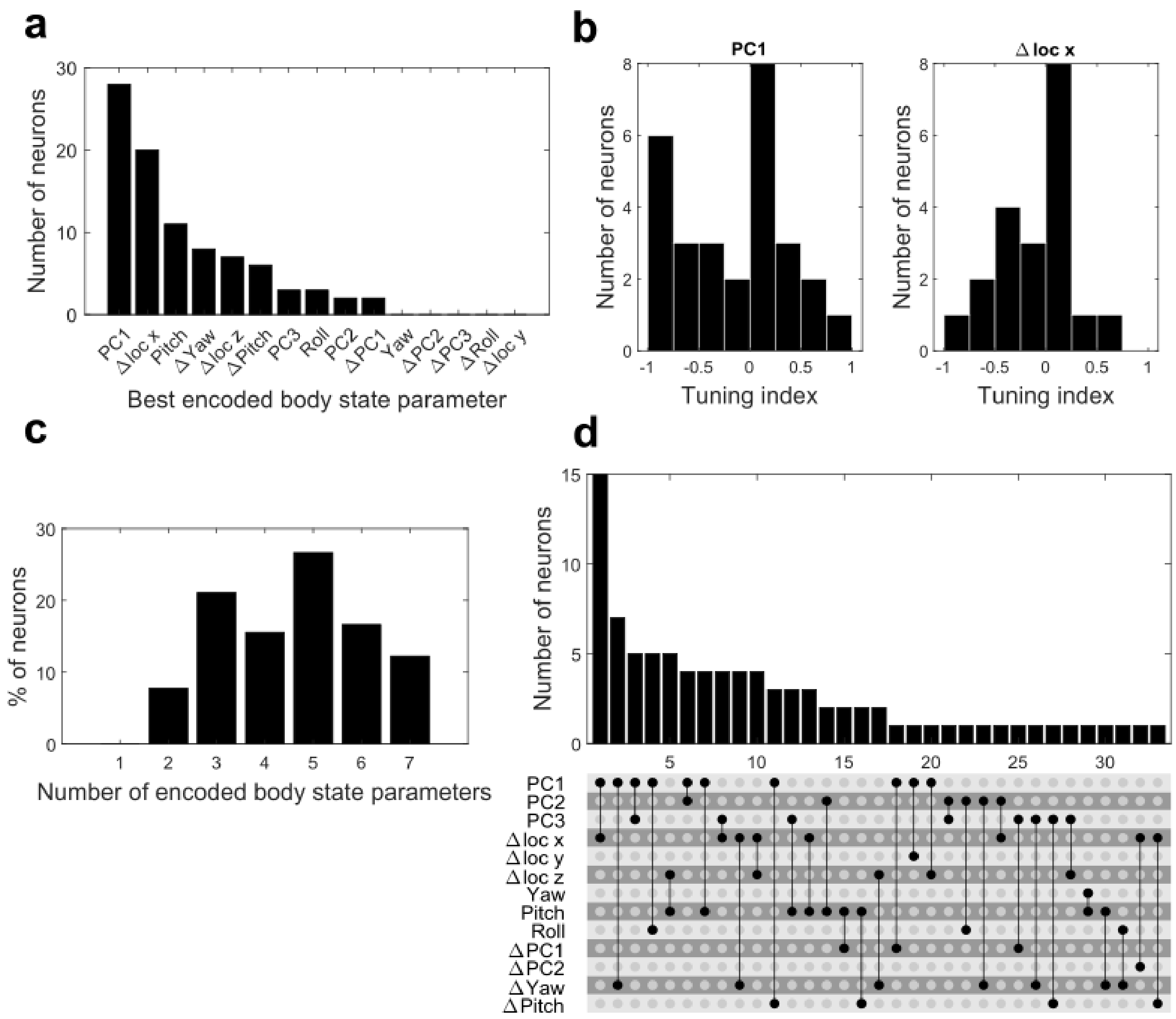


**Supplementary Figure 5. Major aspects of body state encoding observed after ION section are consistent with those observed before ION section. a** Distribution of body state parameters that best predicted neuronal activity ('best predictors'). Same data set as Fig. 4. **b** Histogram of tuning indices to PC1 and Δlocx for neurons where these were the best-predictors (evaluated as in Fig. 5b). **c** Distribution of how many body state parameters each neuron encodes (evaluated as in Fig. 5c). **d** Distribution of neuron types based on the pair of body state parameters that best predict their response. (Top) Frequency of observed neuron types. (Bottom) Combinations of parameters that define a neuron type.
